# Does the Brain’s E:I Balance Really Shape Long-Range Temporal Correlations? Lessons Learned from 3T MRI

**DOI:** 10.1101/2025.03.28.645973

**Authors:** Lydia Sochan, Jessica Archibald, Alexander Mark Weber

## Abstract

A 3T multimodal MRI study of healthy adults (n=19; 10 female; 21.3 - 53.4 years) was performed to investigate the relationship between fMRI long-range temporal correlations and excitatory/inhibitory balance. The study objective was to determine if the Hurst exponent (H) — an estimate of the self-correlation and signal complexity — of the blood-oxygen-level-dependent signal is correlated with the excitatory-inhibitory (E:I) ratio. E:I has been proposed to serve as a control parameter for brain criticality — the theory that the brain operates near a critical point between order and disorder, optimizing information processing and adaptability — which H is believed to be a measure of. Thus, understanding if H and E:I are correlated would clarify this relationship. Moreover, findings in this domain have implications for neurological and neuropsychiatric conditions with disrupted E:I balance, such as autism spectrum disorder, schizophrenia, and Alzheimer’s disease. From a practical perspective, H is easier to accurately measure than E:I ratio at 3T MRI. If H can serve as a proxy for E:I, it may serve as a more practical clinical biomarker for this imbalance and for neuroscience research in general. The study collected functional MRI and magnetic resonance spectroscopy data during rest and movie-watching. H was found to increase with movie-watching compared to rest, while E:I (glutamate/GABA) did not change between conditions. H and E:I were not correlated during either movie-watching or rest. This study represents the first attempt to investigate this connection *in vivo* in humans. We conclude that, at 3T and with our particular methodologies, no association was found. We end with lessons learned and suggestions for future research.

## 1 Introduction

The brain is composed of individual neurons, each computationally-limited, that ultimately work together to produce astonishingly complex operations. Understanding how these neurons produce this emergent behaviour remains one of the great scientific challenges of our time. Recent advances in neuroscience have led to the emergence of the Critical Brain Hypothesis, which posits that the brain operates at a critical point, a state where order and disorder are balanced to enable the most efficient information processing^1–14^. When in a critical state, the brain is optimally responsive to both internal and external stimuli while maintaining a balance between stability and instability^1,2,13,14^. A fundamental theorem proposing how neural networks achieve this delicate balance centers around the excitation-inhibition (E:I) ratio: the balance of excitatory and inhibitory neural activity, often operationalized as the ratio of the primary excitatory and inhibitory neurotransmitters (i.e. glutamate (Glu) and γ-aminobutyric acid (GABA)^6,8–12)^.

*In vivo* measurement of brain criticality can be achieved using several available methods, as systems near a critical state often exhibit fractal-like fluctuations or scale-invariance^15^. Scale-invariance has been observed extensively across various neuronal spatio-temporal scales, from dendritic branching structures^16^ in the spatial domain, to neurotransmitter release^17^, neuronal firing rates^18^, local field potentials^19^, electroencephalography (EEG)^20^ and functional magnetic resonance imaging (fMRI), which measures the blood oxygen level-dependent (BOLD) signal^21–23^ in the time domain. This fractal structure can be quantified by assessing the signal’s long-range temporal correlations (LRTC), which reflect the persistence or memory in a time series, where future values are statistically related to past values over long extended periods. The Hurst exponent (H) is a useful measure for evaluating LRTC^23,24^. An H value between 0.5 and 1 indicates a long-range positive correlation, with large values being followed by large values and small values by small ones. Conversely, an H exponent value between 0 and 0.5 indicates a long-range negative correlation, where large values are likely to be followed by small ones and vice versa. An H value of 0.5 is uncorrelated random noise^25^.

H has emerged as a valuable tool in neuroscience and clinical research. Typically, H values reported in adult brains are above 0.5, with higher H values in grey matter than white matter or cerebrospinal fluid^26,27^. Some key findings from neuroscience research include: a decrease in H during task performance^28,29^; an increase in H during movie watching in the visual resting state network^30^; negative correlations with task novelty and difficulty^31^; increases with age in the frontal and parietal lobes^26^, and hippocampus^32^; decreases with age in the insula, and limbic, occipital and temporal lobes^26^; and H < 0.5 in preterm infants^33^. For a more in depth review, see^23^. In terms of clinical findings, abnormal H values have been identified in Alzheimer’s disease (AD)^34,35^, autism spectrum disorder (ASD)^36–39^, mild traumatic brain injury^40^, major depressive disorder^41,42^ and schizophrenia^39,43^. Crucially, these same disorders have been associated with imbalances in E:I (AD^44,45^; ASD^10,46,47^; depression^48^; and schizophrenia^49^)

In addition to its implications for the critical brain hypothesis, establishing a connection between E:I and H could facilitate simpler estimation of excitatory and inhibitory neurotransmitters, as precise measurement of E:I is technically challenging^50^. Currently, magnetic resonance spectroscopy (MRS) is the only non-invasive method for *in vivo* assessment of the Glu/GABA ratio (excitatory to inhibitory neurotransmitters) in humans^51,52^. Unfortunately, MRS suffers from limited spatial and temporal resolution, with single-voxel spectrosopy sizes between 20 and 27 cm^3^, and scan times between 5-10 minutes^12,50,51^. In addition, quantification of GABA at 3T necessitates the use of a specialized J-difference spectral-editing sequence (such as MEGA-PRESS), in order to isolate GABA (at 3 ppm, 2.3 ppm, and 1.9 ppm) from overlapping peaks (e.g. creatine and glutamate)^53^. The need for this additional sequence therefore reduces the temporal resolution further. If H could function as a substitute for E:I, it could simplify the estimation of E:I in conditions of interest such as ASD, AD, and schizophrenia.

Going beyond clinical observations that imply a link between H and E:I, several studies have attempted to explore this association directly. However, they have primarily relied on computational models or animal studies^6–12^. To date, no *in vivo* study has validated this association in humans. Moreover, the existing research findings have so far been inconsistent, with some reporting positive-linear, negative-linear, or U-shaped relationships between the two variables. Despite these uncertainties, several clinical studies have used fMRI-derived H values as proxies for the E:I ratio^38,39,54^. Therefore, further investigation is needed to establish the true nature of the potential E:I-Hurst relationship and to confirm its presence within the human brain. The current study aimed to explore the potential E:I-Hurst relationship *in vivo*, specifically within the visual cortex during movie-watching and resting periods. We hypothesized that H would increase in the visual cortex during movie watching^30^, and that H would negatively correlate with E:I^11^.

## 2 Methods

### 2.1 Participants

Twenty-seven healthy adult participants were recruited to the study. One participant was not scanned due to claustrophobia while in the scanner. After our analysis and performing quality assurance (see below for details), a further seven participants were removed for having less than ideal MRI data quality, leaving nineteen final participants, between the ages of 21.3 and 53.4 (mean age ± sd: 30.1 ± 8.7 years; 9 males).

### 2.2 Ethics Statement

Written informed consent was obtained from all participants. Ethics approval was granted by the Clinical Research Ethics Board at the University of British Columbia and BC Children’s & Women’s Hospital (H21-02686).

### 2.3 Scanning Procedure

After two anatomical sequences were acquired, participants were instructed to visually fixate on a cross-hair for 24 minutes (see Table 1 and below for more details). During this period, an fMRI, single-voxel semi Localization by Adiabatic SElective Refocusing (sLASER)^55^, and single-voxel MEscher-Garwood Point-REsolved SpectroScopy (MEGA-PRESS)^56^ sequences were acquired (see Figure 1 A and Section 2.4). Next, participants were instructed to watch a nature documentary (Our Planet (2019), Episode 3, “Jungles”^57^) for 24 minutes. During this period, another set of fMRI, sLASER, and MEGA-PRESS sequences were acquired. See Figure 1 B for a visual representation of the scanning protocol. Total scan duration was approximately 1 hour of uninterrupted scanning (i.e. subjects were not removed from the scanner). All participants followed the same order of rest than movie, and all participants saw the same movie segment, beginning at the same time during the scan.

**Figure 1.**
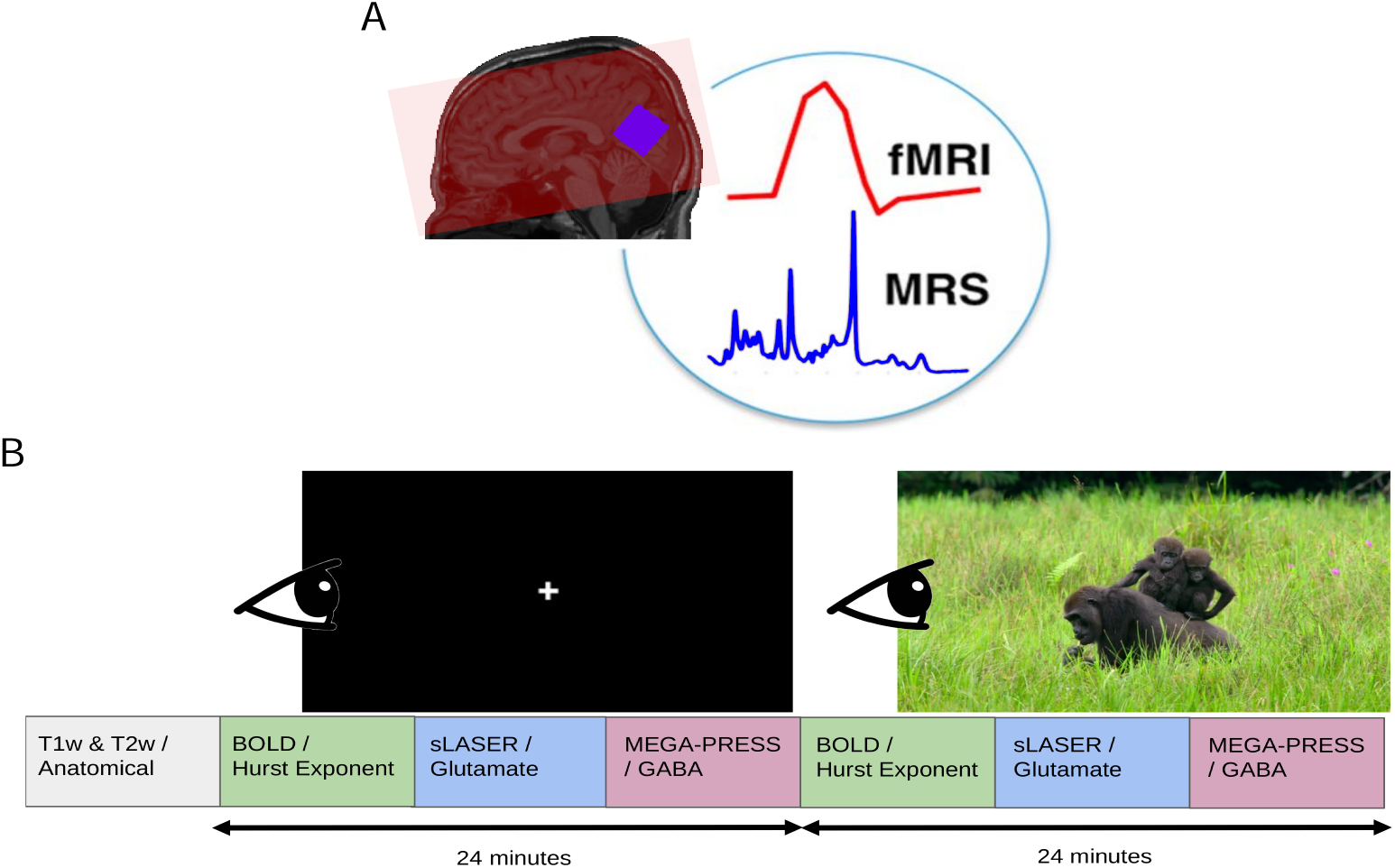
Overview of the MRI acquisition protocol. A) fMRI coverage was across the whole-brain (example coverage in red). MRS voxels were placed in the visual cortex (blue). Background image is of a T1w acquisition from a sample subject. Figure was inspired by Ip et al. (2017)^58^. B) fMRI, sLASER, and MEGA-PRESS were acquired first with participants looking at a white cross for 24 minutes. Next, the same sequences were acquired with the participants viewing a nature documentary for an identical period of time.

**Table 1.**
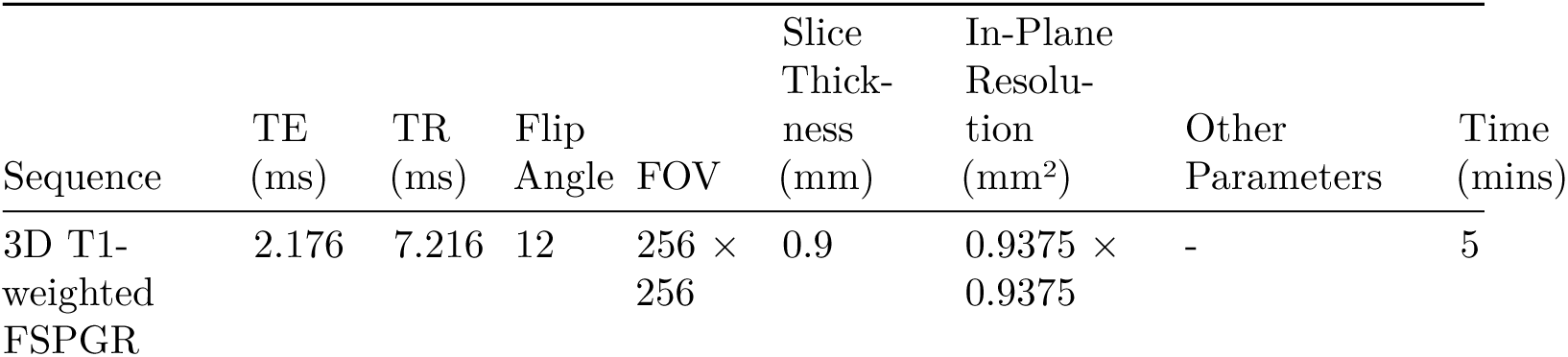
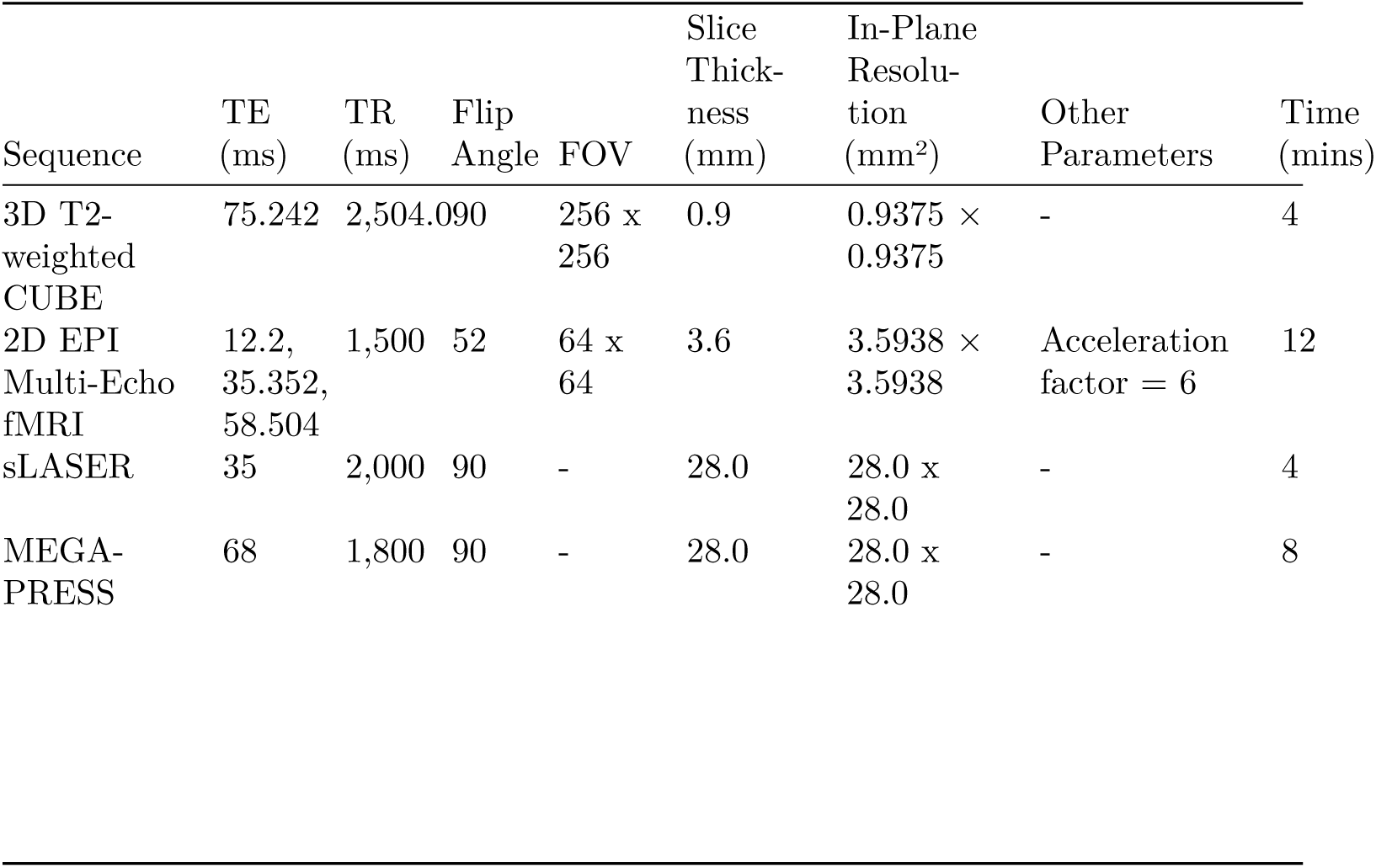
Summary of MRI acquisition details. TE = echo time; TR = repetition time; FOV = field of view; FSPGD = fast spoiled gradient-echo; CUBE = a GE acronym; EPI = echo-planar imaging; fMRI = functional magnetic resonance imaging; sLASER = semi localization by adiabatic selective refocusing; MEGA-PRESS = Mescher-Garwood point-resolved spectroscopy

### 2.4 Acquisition Details

Scans were performed at BC Children’s Hospital MRI Research Facility on a 3.0 Tesla GE Discovery MR750 scanner (scanner software version: DV26.0_R03) with a Nova Medical 32 channel head coil. Participants changed into scrubs and were screened by an MRI technologist. Participants were given wax earplugs and a fiducial was placed on their left temple. Participants were provided with an audio headset and blanket once lying down on the scanner bed. Since visual stimuli were to be rear-projected, position and angle of mirror above patient eyes was adjusted for optimal movie viewing.

The following MRI scans were acquired. A 3D T1-weighted sagittal fast spoiled gradient echo (FSPGR) sequence; a 3D T2-weighted sagittal CUBE; a 2D echo-planar imaging (EPI) multi-echo gradient-echo fMRI sequence; a MEGA-PRESS sequence; and an sLASER sequence. Details are listed in Table 1.

For both MRS sequences, the voxel size was set to 2.8 x 2.8 x 2.8 cm^3^. MRS voxels were rotated and placed in the occipital lobe, aligned along the calcarine fissure. Finally, blip-up and blip-down spin-echo versions of the fMRI sequence were acquired at the end to estimate the B0 non-uniformity map for fMRI phase distortion correction.

### 2.5 Image Processing

Our full image processing pipeline has been can be accessed from our Github account page: github.com/WeberLab/EI_Hurst_Analysis Images were downloaded offline from the scanner in raw Digital Imaging and Communications in Medicine (DICOM) format. DICOM files were then converted to Neuroimaging Informatics Technology Initiative (NIfTI) using Chris Rorden’s dcm2niix^59^ (v1.0.20211006) and then to Brain Imaging Data Structure (BIDS)^60^ format using dcm2bids^61^ (v2.1.6).

#### 2.5.1 Structural Images

The T1w image was corrected for intensity non-uniformity with N4BiasFieldCorrection^62^ and distributed using ANTs^63^ (v2.3.335) to be used as a T1w-reference for the rest of the workflow. The T1w-reference was skull-stripped using a Nipype^64^ implementation of antsBrainExtraction.sh from ANTs; OASIS30ANTs was used as a target template. Fast^65^ (FSL^66^ v.6.0.5.1:57b01774, RRID: SCR_002823) was used for brain tissue segmentation into cerebrospinal fluid (CSF), white matter (WM), and gray matter (GM). Brain surfaces were reconstructed with recon-all^67^ (FreeSurfer^67^ 7.3.2, RRID: SCR_001847). The previously-estimated brain mask was refined with Mindboggle^68^ (RRID:SCR_002438) to reconcile ANTs-derived and FreeSurfer-derived segmentations of cortical GM. AntsRegistration^63^ (ANTs 2.3.3) was used to perform volume-based spatial normalization to two standard spaces: MNI152NLin2009cAsym and MNI152NLin6Asym. Normalization used brain-extracted versions of both T1w reference and T1w template.

#### 2.5.2 fMRI

Using fMRIPrep^69^, the shortest echo of the BOLD run was used to generate a reference volume (both skull-stripped and skull-included). Head-motion parameters with respect to the BOLD reference (transformation matrices as well as six corresponding rotation and translation parameters) were estimated before spatiotemporal filtering using mcflirt^70^ (FSL v6.0.5.1:57b01774). The fieldmap was aligned with rigid registration to the target EPI reference run. Field coefficients were mapped to the reference EPI using the transform. BOLD runs were slice-time corrected to 643 ms (half of slice acquisition range of 0-1290 ms) using 3dTshift from AFNI^71^ (RRIS: SCR_005927). To estimate T2* map from preprocessed EPI echoes, a voxel-wise fitting was performed by fitting the maximum number of echoes with reliable echoes in a particular voxel to a monoexponential signal decay model with nonlinear regression. Initial values were T2*/S0 estimates from a log-linear regression fit. This calculated T2* map was then used to optimally combine preprocessed BOLD across echoes using the method by Posse et al. (1999)^72^. The generated BOLD reference was then co-registered (6 degrees of freedom) to the T1w reference with bbregister (FreeSurfer^67^) using boundary-based registration. First, a reference volume and its skull-stripped equivalent were generated with fMRIPrep. Confounding time series were calculated from preprocessed BOLD: framewise displacement (FD), DVARS, and three region-wise global signals. Tedana^73^ was then used to denoise the data by decomposing the multi-echo BOLD data via principal component analysis (PCA) and independent component analysis (ICA). The resulting components are automatically analyzed to determine whether they are TE-dependent or -independent. TE-dependent components were classified as BOLD, while TE-independent components were classified as non-BOLD and were discarded as part of data cleaning. Participants were excluded from further analysis if their mean FD was > 0.15 mm.

### 2.6 MRS

sLASER data were processed and fit to a spectrum using Osprey^74^ (v2.4.0). Full width half-maximum (FWHM) of the single-Lorentzian fit of the N-acetylaspartate (NAA) peak were calculated for quality assurance purposes. The MRS voxel was co-registered to T1w reference image and segmented by SPM12^75^ into CSF, GM, and WM. Metabolites were water-scaled as well as tissue- and relaxation-corrected by the Gasparovic et al. (2006) method^76^. Glutamate is challenging to capture due to its signal overlaps with other metabolites^77^. In particular, Glu shares a similar chemical structure with glutamine (Gln) which causes the spectral features of Glu to be contaminated with Gln^78^. As a result, we decided to report Glx values, which co-reports Glu and Gln to avoid errors in spectral assignment, especially since it is controversial whether Glu can reliably be separated from Gln at 3T^79,80^.

MEGA-PRESS data were processed and fit with Osprey, and were relaxation-, tissue- and alpha-corrected using the Harris et al. (2015) method^81^. Due to the J-editing sequence of MEGA-PRESS, a challenge of GABA quantification is macromolecule quantification^81^. As a result, we report GABA+, a measure which co-reports GABA with macromolecules. Macromolecules (MM) are expected to account for approximately 45% of the GABA+ signal^81^. While a macromolecule-suppressed estimate of GABA seems ideal, a recent 25-site and multi-vendor study conducted at 3T found that GABA+ showed much lower coefficient of variation than MM-suppressed GABA, meaning that GABA+ is more consistent across research sites and MRI vendors (i.e., Philips, GE, Siemens)^82^. Moreover, GABA+ shows greater reliability for both creatine-referenced and water-suppressed estimates^82,83^. MM-suppressed GABA and GABA+ estimates are also correlated, albeit weakly- to moderately-so^81–83^. Consequently, we report GABA+ to allow for easier comparison of our results to other studies as well as reproducibility.

Basis sets were created using MRSCloud^84,85^. For sLASER, ‘localization’ was set to ‘sLASER’, ‘vendor’ to ‘GE’, ‘editing’ to ‘UnEdited’, and ‘TE’ to 35. For MEGA-PRESS, ‘localization’ was set to ‘PRESS’, ‘editing’ to ‘MEGA’, ‘TE’ to 68, ‘edit on’ to ‘1.9’, ‘edit off’ to ‘7.5’, and ‘pulse duration’ to ‘14’. Metabolites included for both basis sets were: Asc, Asp, Cr, CrCH2, EA, GABA, GPC, GSH, Gln, Glu, Gly, H2O, Lac, NAA, NAAG, PCh, PCr, PE, Ser, Tau, mI, and sI. Excitatory-inhibitory ratio (E:I) was calculated as [Glx from sLASER in i.u.]/[GABA+ from MEGA-PRESS in i.u.], a common practice to report E:I using MRS^86^.

Participants were excluded from further analysis if any of their MRS scans had FWHM > 10. Originally we intended to calculate, report and analyze MRS as tissue-corrected quantitative values. However, the Osprey software was reporting Glx metabolite concentrations from the sLASER sequence that were an order of magnitude larger than reported values in the literature (i.e. x10). Attempts to re-analyze the sLASER data using fsl_mrs^87^ resulted in similarly large concentrations. Despite several attempts, we were unable to locate the source of this error. Therefore we decided to analyze and report metabolite to creatine (tCr) ratios (i.e. Glx / tCr and GABA+ / tCr), which agreed well with reported literature values^88–92^. In order to be confident in our results, however, we re-ran all MRS analyses, acquiring quantitative Glx and GABA+ values from the MEGA-PRESS sequence only using the software Gannet^93^ (v3.3.0). Values were relaxation-, tissue- and alpha-corrected using the Harris et al. (2015) method^81^. Results using Glx and GABA+ from the Gannet analysis of MEGA-PRESS only are reported in the Supplementary Material.

### 2.7 Hurst Exponent Calculation

Hurst exponent was calculated from the power spectrum density (PSD) of the BOLD signal. A log-log plot was used, where log power was plotted against log frequency; generally, if a log-log plot results in a linear relationship, it is assumed that the mean slope of this line represents the power-law exponent^5^. A PSD shows the distribution of signal variance (‘power’) across frequencies. Complex signals are classified into two categories: fractional Gaussian noise (fGn) and fractional Brownian motion (fBm)^24,94^. The former is a stationary signal (i.e., does not vary over time), while the latter is non-stationary with stationary increments^24^. Most physiological signals consist of fBm, but fMRI BOLD is typically conceptualized as fGn once motion-corrected; otherwise put, unprocessed BOLD signal is initially fBm which is converted to fGn with appropriate processing^95^. fBm and fGn require distinct H calculation methods^24^. PSD was estimated using Welch’s method^96^ from the Python Scipy.Signal library^97^. Data were divided into 8 windows of 50% overlap and averaged. The spectral index, β, was calculated from the full frequency spectrum. The spectral index was then converted to H using the following equation^24,98^:

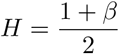

Since it cannot be assumed that all fBm is removed from the signal, we employed the ‘extended Hurst’ (H’) concept in this study: for 0 < H < 1, the signal is understood as fGn, while for 1 < H < 2, the signal is understood to be fBm^30,99,100^. More generally, it is assumed that when 0.5 < H < 1.5, the signal displays 1/f behaviour^5^. H was calculated for all voxels in the brain of each subject. A brain mask was then applied which included only GM and the region of the MRS voxel in the visual cortex. H was averaged across the brain mask area, using only non-zero voxels.

### 2.8 Statistics

All statistical analyses were performed using R^101^ and RStudio (v2023.06.0+421). Difference of means of H, Glx, and GABA+ between rest and movie conditions were calculated using paired Student’s t-tests^102^. Correlations between H and Glx, GABA+ and E:I were calculated using Pearson’s method^103^. Finally, a linear mixed-effect model was used to test for the correlation of H and E:I while controlling for possible confounding effects of MRS FWHM and fMRI motion (see Results).

## 3 Results

### 3.1 Participant Demographics

Twenty-seven participants were originally recruited for the study. Twenty-six of these participants were successfully scanned, but one participant experienced claustrophobia and chose not to continue. Of the remaining 26 participants, 19 were included in the final analysis: two were removed due to low MRS quality (FWHM > 10) and five were removed due to low fMRI quality (mean FD > 0.15 mm). See Figure 2. The final study sample included 9 males and 10 females between ages 21.3 and 53.4, with a mean age and standard deviation of 30.1 ± 8.7 years.

**Figure 2.**
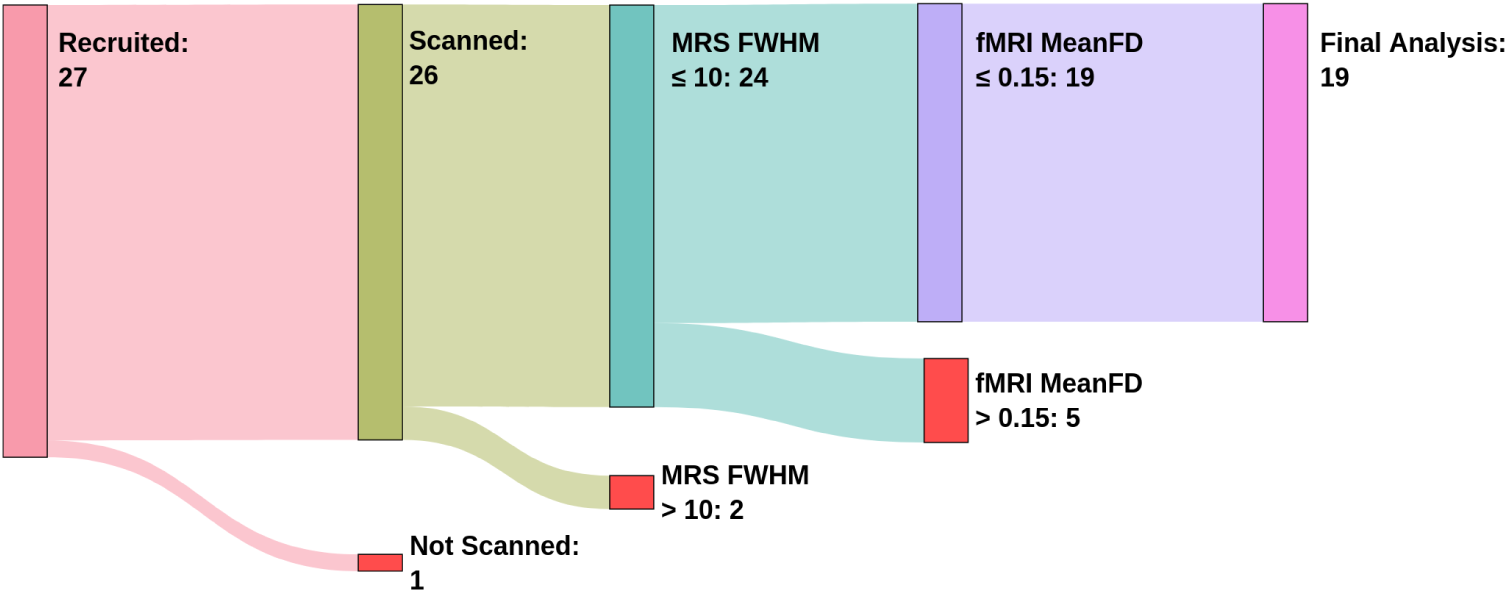
Flowchart of participant, recruitment, scanning and exclusion. The number of participants at each stage is indicated within each node. Participants were excluded based on MRS FWHM and fMRI Mean FD thresholds, resulting in 19 participants included in the final analysis

### 3.2 Data Quality

After exclusion, FWHM (mean ± sd) at rest in sLASER and MEGA-PRESS were 8.52 ± 0.7 and 7.02 ± 0.7, respectively. During movie watching, FWHM in sLASER and MEGA-PRESS were 8.37 ± 0.65 and 6.98 ± 0.86. Glx and GABA+ were tested for associations with FWHM values during rest and movie watching. Glx during rest was found to be negatively correlated with FWHM (r = −0.53, p = 0.02). No other correlations were significant. An average of all MRS voxel placements can be seen in Figure 3 A, and a sample of the Osprey sLASER and MEGA-PRESS spectrum fits at rest can be seen in Figure 3 B and C, respectively.

**Figure 3.**
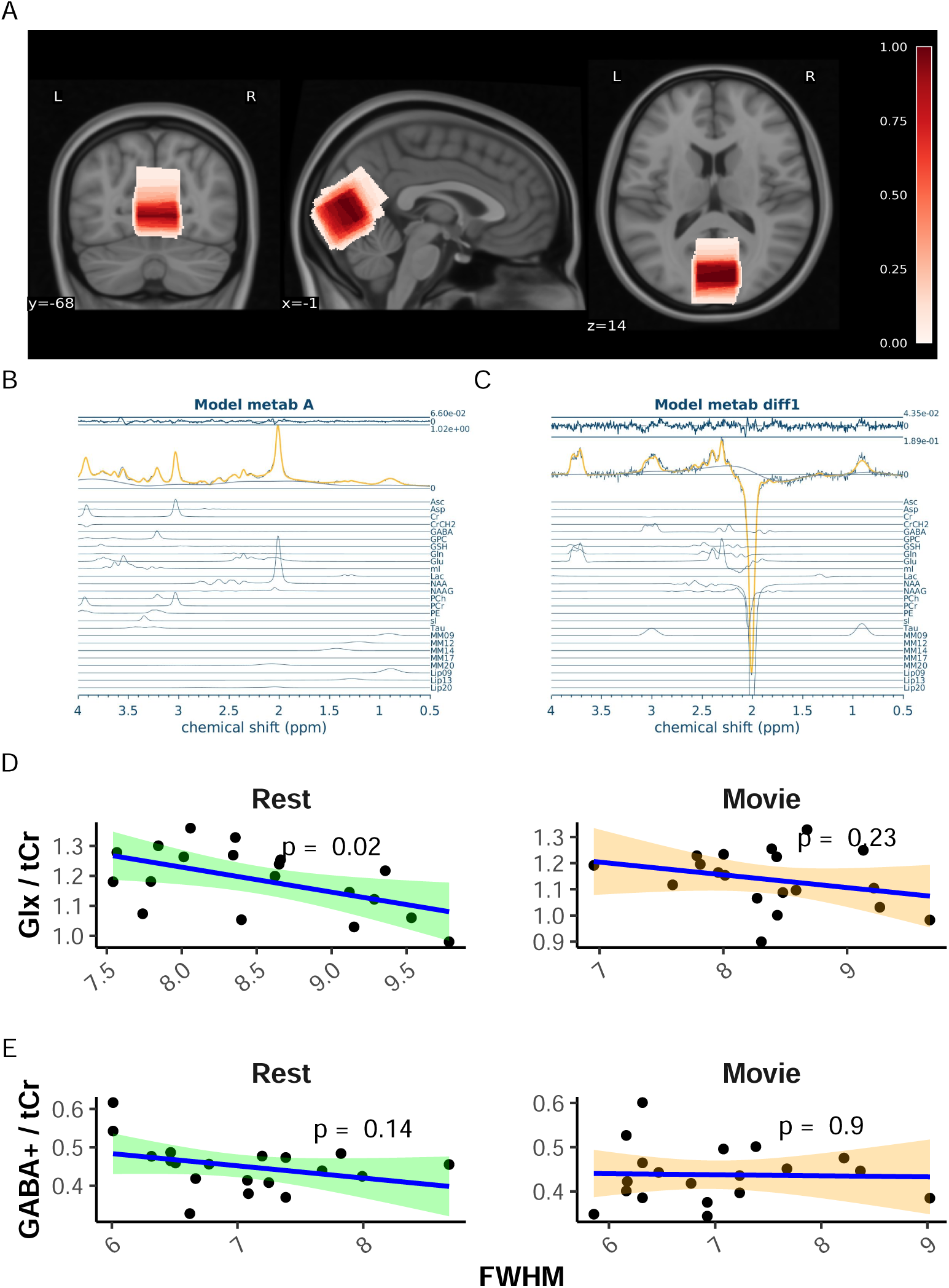
MRS data quality. A) Average MRS location. Overlay in red of the average voxel location across all nineteen participants, from 0 (white) to 1 (dark red). Darker red represents voxel locations shared by all participants, while white represents voxel locations unique to participants. MRS voxels were registered to MNI A sample of the combined grey-matter and MRS voxel mask used to average H values, along with a sample Hurst exponent map, and sample fits for H calculation during rest can be found in Figure 4 A, B, and C, respectively. Mean FD was not correlated with H during rest (r = −0.33, p = 0.16 but was moderately negatively correlated with H during movie watching (r = −0.50, p = 0.03; see Figure 4 D).

**Figure 4.**
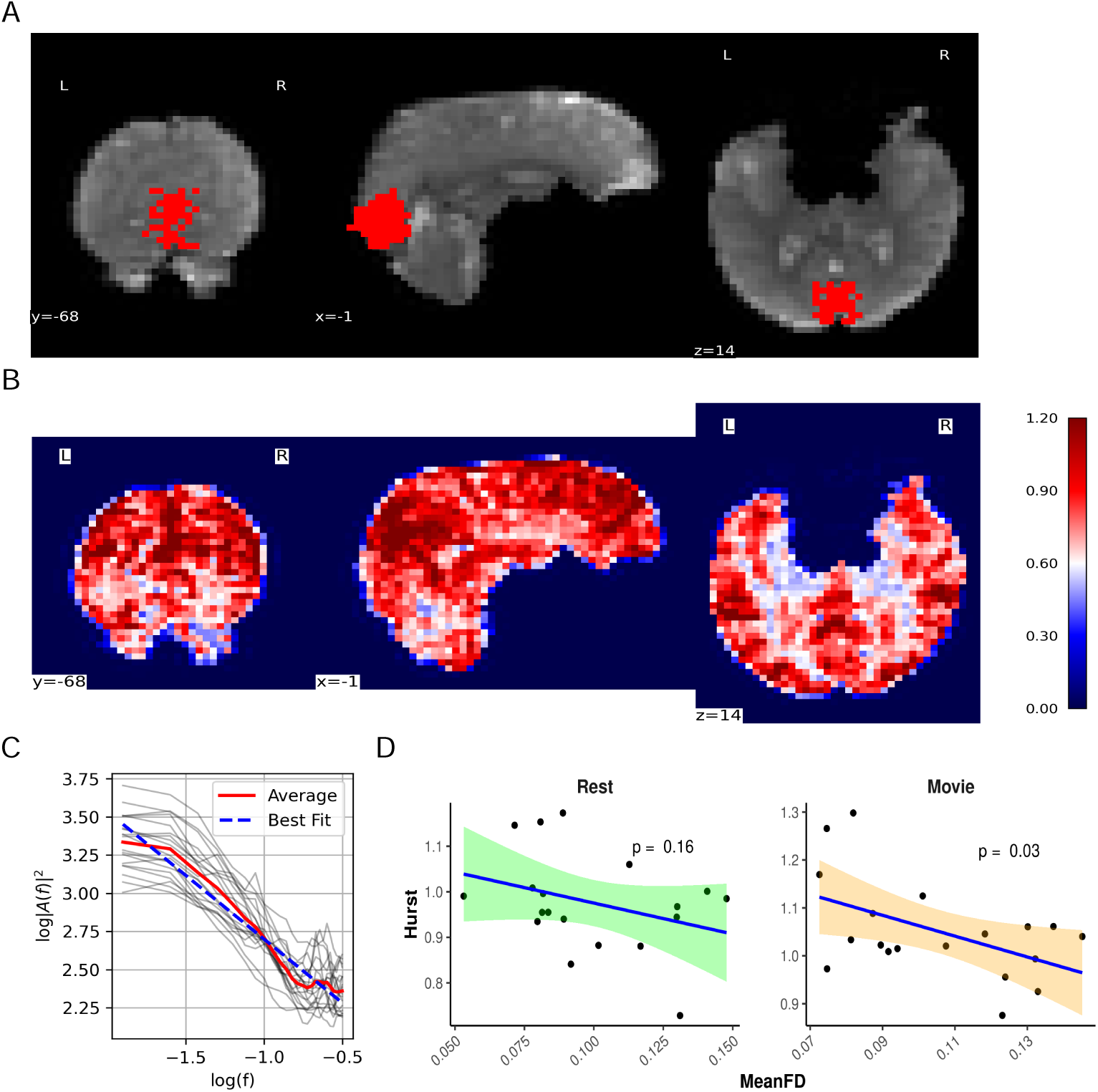
fMRI data quality. A) sample fMRI grey-matter mask within the MRS voxel. Background: a sample coronal, sagittal, and axial slice is displayed of the mean fMRI scan from the rest acquisition. Foreground: the greymatter/MRS mask used to calculate mean H. B) Sample H map for whole brain. Coronal, sagittal, and axial views of are shown, colour-coded by H values. Colour-coding shows evident tissue differentiation by H value (i.e., gray matter (red), white matter (light red), and cerebrospinal fluid (white)). C) Sample PSD spectrum. All participants’ PSD spectrums during rest are plotted using light grey lines. Mean PSD is plotted in solid red Mean linear regression line is plotted in dashed blue. H for each participant was calculated from the slope of their mean linear regression line. D) Correlation plots of H vs. mean FD for rest (green) and movie (orange).

### 3.3 H and E:I

Mean ± sd of metabolites, E:I, and H during rest and movie are reported in Table 2. Boxplots between rest and movie can be seen in Figure 5. Neither Glx nor GABA+ were different between movie and rest conditions. E:I ratio did not change between conditions either. H was found to be greater during movie watching than rest.

**Table 2.** Summary of main results.

|  | Rest | Movie | p-value |
| --- | --- | --- | --- |
| **Glx / tCr** | $1.19 \pm 0.11$ | $1.14 \pm 0.11$ | 0.09 |
| **GABA+ / tCr** | $0.45 \pm 0.06$ | $0.44 \pm 0.06$ | 0.50 |
| **E:I Ratio** | $2.68 \pm 0.49$ | $2.65 \pm 0.45$ | 0.82 |
| **H** | $0.98 \pm 0.98$ | $1.05 \pm 1.05$ | $< 0.01$ |

**Figure 5.**
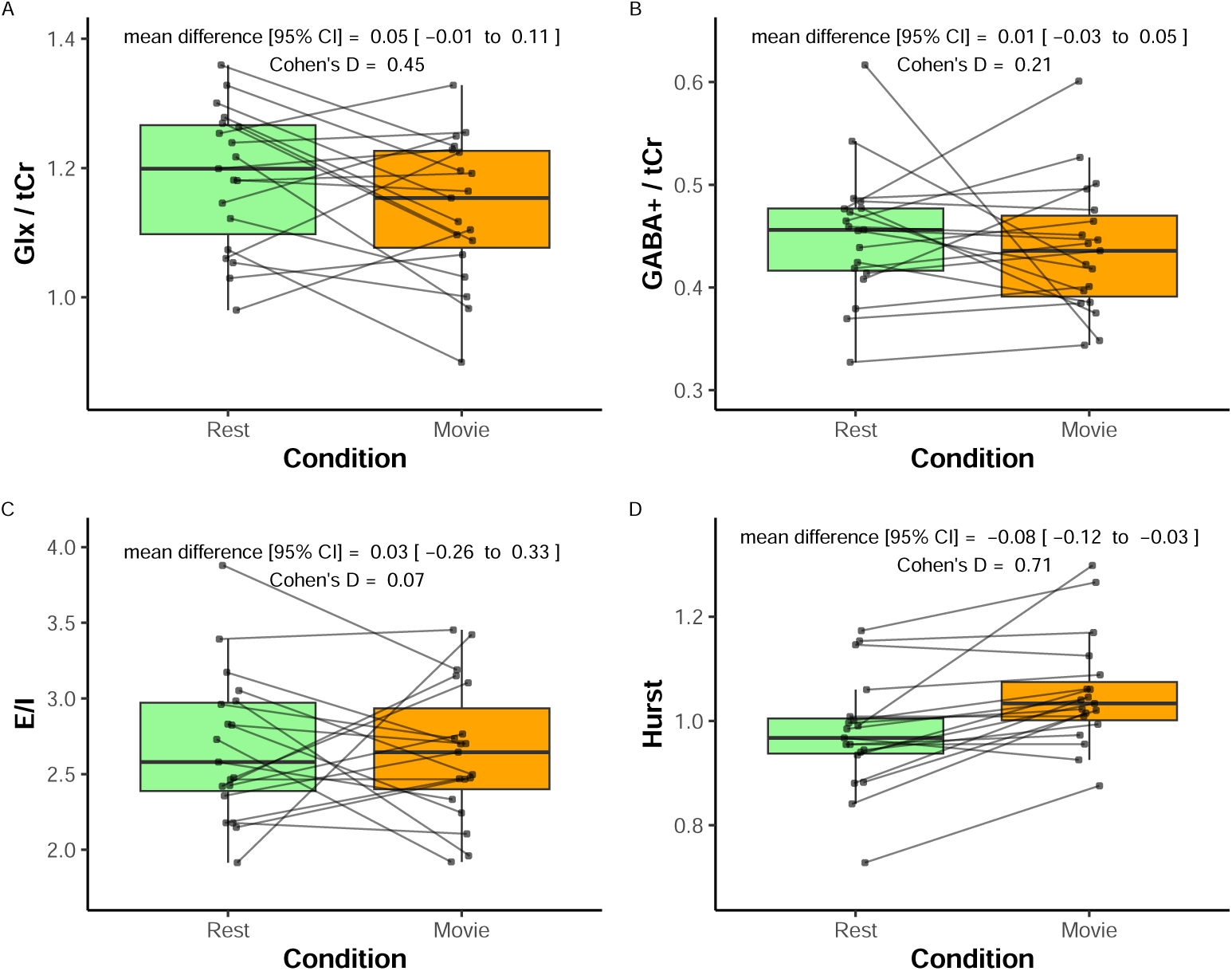
Paired comparison of metabolite values during rest (green) and movie (orange) conditions. A) Glx / tCr; B) GABA+ / tCr; and C) E:I. Paired dots represent the same participant across conditions. Mean difference with 95% confidence intervals, and as Cohen’s D are reported at the top of each plot.

H was not found to correlate with Glx, GABA+, or E:I, during rest or movie (Figure 6).

**Figure 6.**
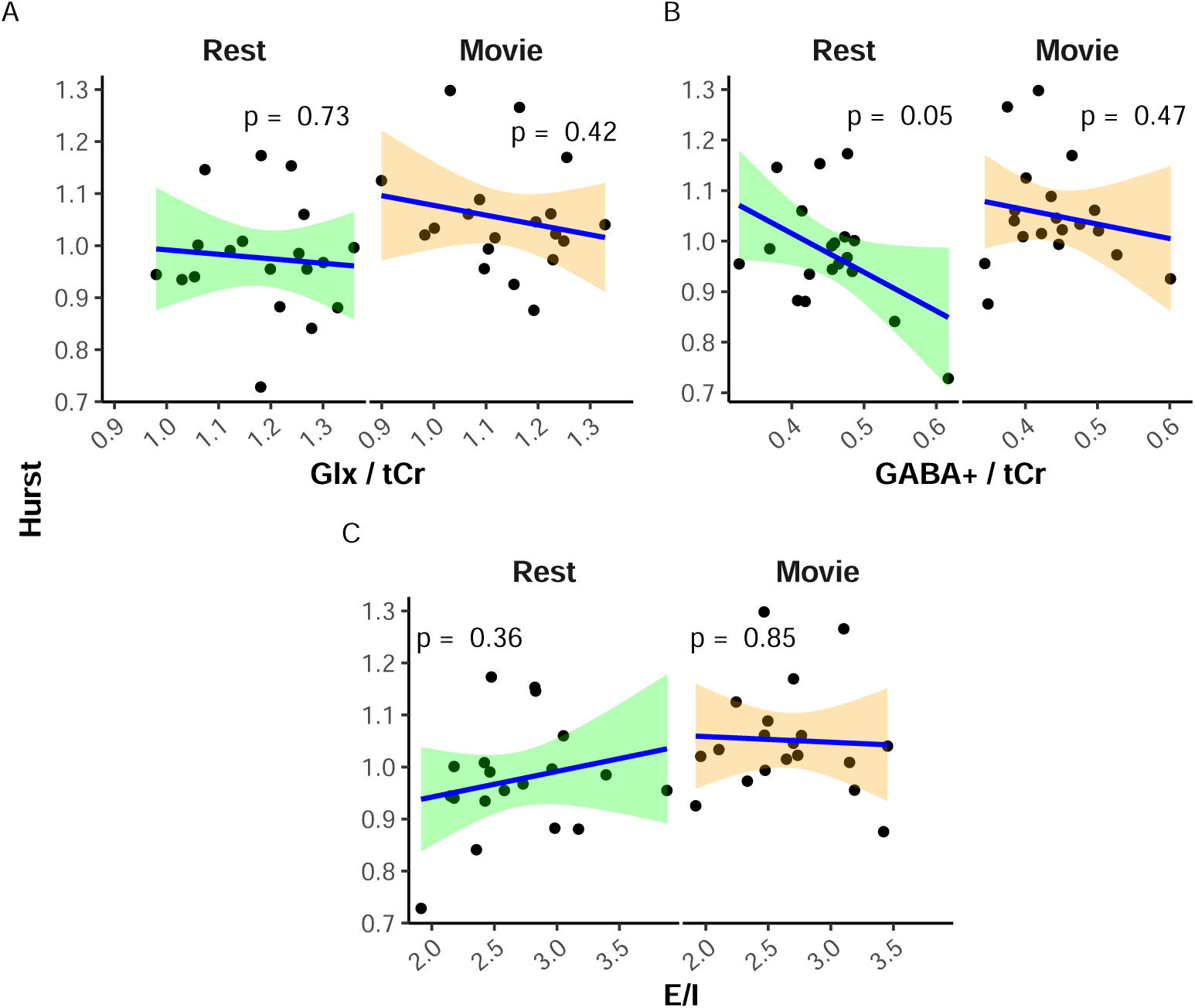
Scatter plots of H vs. metabolites. A) Glx / tCr; B) GABA+ / tCr; and C) E:I. Rest is in green, while movie is in orange. p-values are reported at the top of each plot.

To ensure that mean FD and FWHM were not confounding our results, we also ran a linear mixed-effects model with H as dependent variable, Glx/GABA+ as primary predictor of interest, and mean FD, FWHM(Glx), and FWHM(GABA+) as covariates, Condition (rest vs. movie) as fixed effects, and subjects as random effects:

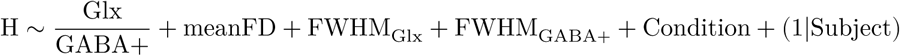

The model’s total explanatory power was substantial (conditional r^2^ = 0.69), with fixed effects alone (marginal r^2^) equal to 0.19. The model’s intercept, corresponding to EI = 0, Condition = Movie, FWHMs_Glx_ = 0, FWHM_GABA+_ = 0 and meanFD = 0, was 0.98 (95% CI [0.41, 1.56], t(30) = 3.34, p = 0.002). Within this model, only the effect of Condition [Rest] was statistically significant (beta = −0.09, 95% CI [-0.13, −0.04], t(30) = −3.99, p < 0.01).

### 3.4 Re-analysis using Gannet

Re-running our analysis using Glx and GABA+ values from Gannet did not result in any significant changes from the results reported above. We did, however, obtain quantitative Glx values much closer to what has been reported in the literature. We have included these results in the Supplementary Material.

## 4 Discussion

We report here the first *in vivo* human study of the E:I-H relationship. An increase in H was observed during movie-watching as compared to rest, indicating stronger long-range temporal dependencies in BOLD activity during visual stimulation. No difference was found in Glx, GABA+, nor Glx/GABA+ between rest and movie-watching. Furthermore, no association was found between H and Glx, GABA+, nor Glx/GABA+. Attempting to control for mean FD and FWHM using a linear mixed-effects model did not alter these results.

Our finding that H increases during movie-watching compared to rest is consistent with a previous study from our lab^30^, which reported increased H in the visual network (from Yeo’s seven resting-state networks^104^) during movie-watching using data from the Human Connectome Project^105^. While this increase in H is consistent with our prior findings, other studies have reported decreases in H during active tasks^28,29,31,106^. Our results suggest that the naturalistic, passive nature of movie-watching elicits a different effect on H compared to more active tasks. This is consistent with literature indicating distinct neural responses and BOLD signal characteristics between conventional active visual tasks and naturalistic passive visual stimuli^30,107^. Richer scaling properties (higher H) during movie-watching may support the continuous perception of visual stimuli^30^.

We found no changes in either Glx or GABA+ between conditions. Given that no difference was observed for either metabolite, it is unsurprising that E:I did not change either. This was unfortunate as our study was designed to include two conditions in order to elicit changes in H, Glx, and GABA+, aiming to confirm that our experiment induced measurable alterations in these metrics. Establishing such changes would have provided a foundation for directly examining the correlation between H and the E/I ratio. However, because Glx and GABA+ did not show significant changes between conditions, our main results are likely to be met with doubt. Without evidence of condition-induced variations in Glx and GABA+, it remains unclear whether the absence of correlation reflects a true lack of association or insufficient sensitivity of our measurement approach to detect changes in these metabolites.

A recent meta-analysis of Glu/Glx studies^77^ reported a minimal task-induced increase in Glu/Glx within the visual cortex; however, this finding was not linked to any visual stimuli task, but rather to pain, learning, and motor tasks. Additionally, many of the studies included in the review were conducted at 7T, offering increased sensitivity for detecting changes in Glu/Glx. Nonetheless, several studies have demonstrated changes in Glu/Glx at 3T (e.g.,^108–110)^. Regarding our GABA+ findings, the same metaanalysis^77^ reported no task-dependent change in GABA within the visual cortex. This may be attributable to technical difficulties associated with capturing GABA levels using MRS at 3T due to its low concentration and signal overlap with more abundant metabolites^77^. Nevertheless, it is worth noting that several studies have successfully reported changes in GABA at 3T using different paradigms and/or regions of interest (e.g.,^111–113)^.

Another reason we may not have found changes in Glx or GABA+ may be due to use of a block-design: collecting 24 minutes of rest data, then collecting 24 minutes of movie-watching data. While a block design has the advantage of a more robust metabolite quantification due to greater signal averaging, brain homeostatis during these long blocks may lead to an erasure of any real metabolic changes^110,114^. Indeed, Pasanta et al. (2023)^77^ found the magnitude of effect sizes were observed to be smaller for block designs than event-related designs.

However, it may be possible that our results do support the idea that H and E:I are not directly linked; at least not in a simple manner. This is would perhaps not be surprising given the large disparity of findings in the literature, especially with regard to the directionality and linearity of the proposed E:I-Hurst relationship^6–12^ (see Table 3). The heterogeneity across these E:I-Hurst studies highlights the challenges of studying this phenomenon as well as the complexity of any potential relationship between these two metrics. It may be that, given the mixed-findings of these studies in combination with our data, an E:I-Hurst relationship — should it exist — may depend in part on how the data is collected. This could include variables such as the experimental setup, sampling methods, or data analysis techniques used. Further research is needed to elucidate the true nature of any potential E:I-Hurst relationship and better understand the complexities involved in studying this phenomenon.

**Table 3.** Summary of methods for existing E:I-Hurst studies.

| Citation | Study Type | H Data Type | H Calculation Method | E:I Type | E:I-Hurst Relationship |
| --- | --- | --- | --- | --- | --- |
| Poil et al. (2012)<sup>7</sup> | Computational with in-house simulated model | Neuronal avalanche size | Detrended fluctuation analysis (DFA) | Structural: number of E-to-I neurons | Inverse U |
| Bruining et al. (2020)<sup>10</sup> | Computational with model by Poil et al. (2012); modified in-house | Neuronal oscillation amplitude | DFA | Structural: number of E-to-I synapses | Inverse U |
| Gao et al. (2017)<sup>12</sup> | Computational in vivo in rats and macaques | Local field potential (LFP) | PSD | Estimated from LFP | Positive linear |
| Lombardi et al. (2017)<sup>8</sup> | Computational with in-house model | Neuronal avalanche size | PSD | Structural: number of E-to-I neurons | Negative linear |
| Trakoshis et al. (2020)<sup>11</sup> | Computational with simulated data; in vivo in mice | fMRI BOLD signal | Wavelet-based maximum likelihood method | E-to-I synaptic conductance | Positive linear |

Finally, it is also possible that while an E:I-Hurst relationship exists, it is not observed within the visual cortex. This theory seems plausible given that MRS studies of disrupted E:I, mostly conducted within the context of adult autism spectrum disorder, have found changes in E:I within other brain regions such as the anterior cingulate cortex, frontal lobe, or temporal lobe^50^. Moreover, findings with reference to changes in excitatory or inhibitory neurotransmitters within the visual cortex tend to be difficult to capture, perhaps indicating that E:I shows less changes in this region^77^. However, this would suggest that the E:I-H relationship would be region dependent, and therefore not a generalizable theory as it is often portrayed.

### 4.1 Limitations and Strengths

Beyond the limitations already mentioned (field strength, visual cortex, passive task, block design), another limitation was our small sample size. As this was a pilot study, 26 participants were initially scanned. Once individuals were excluded for poor MRI quality, only 19 participants’ data were analyzed. With this small sample size, it is difficult to make conclusions about a concept as complex as the E:I-Hurst relationship. We hope that by publishing our work, future researchers can use our reported effect sizes to calculate potential sample sizes. We would also like to list some of the strengths of our study, which include: using sLASER (as opposed to PRESS), which has been shown to have enhanced detection of complex multiplets such as Glu^115^; using the J-editing sequence MEGA-PRESS for improved GABA detection^53^; using a large MRS voxel size (∼22 ml) as per consensus recommendations^116–118;^ measuring H within the same region as our single-voxel MRS; and using multi-echo fMRI for improved motion artifact regression^119^.

### 4.2 Lessons for Future Researchers

Finally, we hope that by publishing our findings, we can provide some guidance to future researchers. The following is a non-exhaustive list of suggestions for future work:

- future studies should consider using ultra-high-field 7T MRI, which provides improved spectral resolution and more reliable detection of Glu and GABA compared to conventional 3T MRI;
- examining paradigms that have consistently demonstrated alterations in these metabolites, such as pain studies, may enhance the likelihood of detecting changes in Glu and GABA^108,109,120^.
- exploring brain regions beyond the visual cortex, such as the anterior cingulate cortex, which is commonly implicated in pain processing^120–123;^
- including a more diverse participant sample — beyond healthy controls — may help capture a wider range of H and E:I values, potentially improving sensitivity to metabolite-related changes;
- increasing sample size in order to detect small effect sizes;
- using an event-related design, as opposed to the block design we used, may provide more sensitivity to detecting rapid and transient neural responses due to its ability to isolate specific events within the scan session^124^. However, see Pasanta et al. (2023)^77^ for a longer discussion and possible downsides to this approach;
- and finally, using a combined fMRI-MRS sequence^58,89^ to measure BOLD and Glu/GABA near-simultaneously.

Together, these considerations may help in overcoming the limitations observed in the present study and contribute to a clearer understanding of the potential relationship between E:I and H.

## 5 Conclusion

In conclusion, our findings do not support a relationship between H and E:I in the visual cortex either during rest or during movie-watching at 3T in humans. In addition, while we found a task-related change in H, we did not find any changes in Glu, GABA, or E:I between movie and rest. Comparing our findings to the broader literature, E:I balance may be too subtle to be detected with conventional 3T MRS methods. With regards to the broader E:I-Hurst relationship, we similarly suggest that either this relationship is insufficiently captured with our methods, or that the relationship between these two variables may be more complex than originally envisaged — perhaps they are not directly related, but rather connected through other mediating variables in a non-linear fashion. To our knowledge, this is the first *in vivo* human study to test for this relationship. It is our hope that as the literature grows, more authors will examine this relationship with respect to other brain regions and using other methods, and will use the lessons learned in this study to inform their own. Hopefully then it will be possible to corroborate findings to probe the complex relationships that may exist with regards to H and E:I in the human brain.

## Data availability statement

All code used in this paper is available at github.com/WeberLab/EI_Hurst_Analysis. The raw MRI data used in this paper is available by contacting the author. The manuscript was written in a ‘reproducible manner’: the entire manuscript, including statistics reported, figures, and tables, can be reproduced here: weber-lab.github.io/EI_Hurst_Manuscript/

## Competing interests

The authors declare they have no known competing interests.

## Author contributions

LS performed data curation, formal analysis, investigation, software, visualization, and wrote the original draft. JS helped with validation, visualization, and reviewed and edited the manuscript. AMW was responsible for conceptualization, data curation, formal analysis, funding acquisition, investigation, methodology, project administration, resources, supervision, validation, visualization, and writing - review & editing.

## Acknowledgements

The work presented in this paper was supported in part from funding from the British Columbia Children’s Hospital Foundation, and an NSERC Discovery Grant.

## Supplementary Material

### Gannet Results

An example fit result from Gannet of a subject at rest is shown in Figure 7. Mean ± sd of metabolites and E:I when analyzed using Gannet, during rest and movie, are reported in Table 4. Boxplots between rest and movie can be seen in Figure 8. Neither Glx nor GABA+ were different between movie and rest conditions. E:I ratio did not change between conditions either.

**Table 4.** Summary of main metabolite results using Gannet.

|  | Rest | Movie | p-value |
| --- | --- | --- | --- |
| **Glx** | 9.32 ± 0.85 | 9.19 ± 0.6 | 0.30 |
| **GABA+** | 3.09 ± 0.32 | 3.03 ± 0.33 | 0.22 |
| **E:I Ratio** | 3.04 ± 0.3 | 3.06 ± 0.26 | 0.68 |

**Figure 7.**
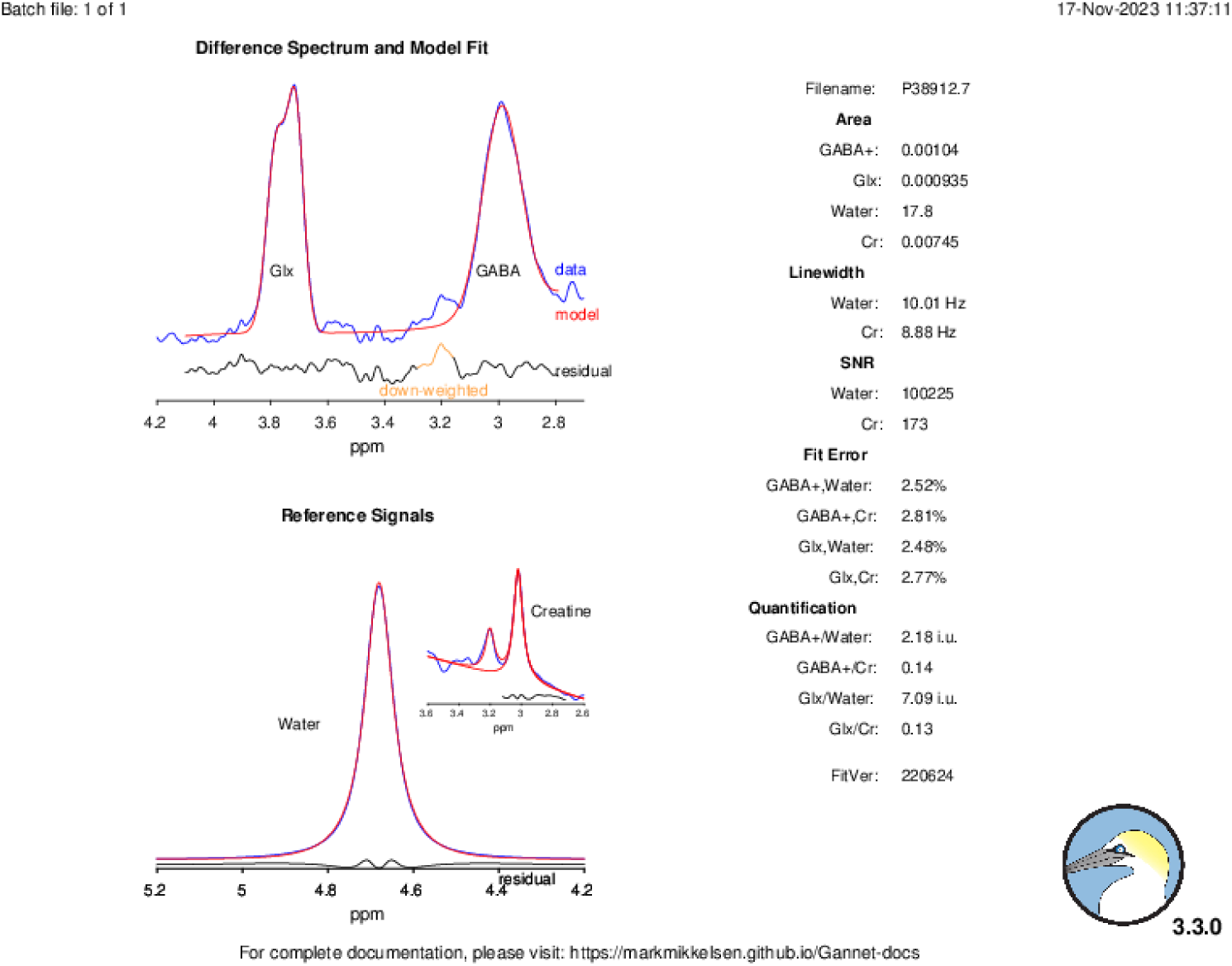
Gannet Fit. Sample Gannet fit result from a subject a rest. Glx (3.75ppm) and GABA (3) data (blue) and model fit (red) are shown top left. Water and creatine reference signals are shown bottom left. Table on right shows GABA+ and Glx values before tissue and tissue- and alpha-correction.

**Figure 8.**
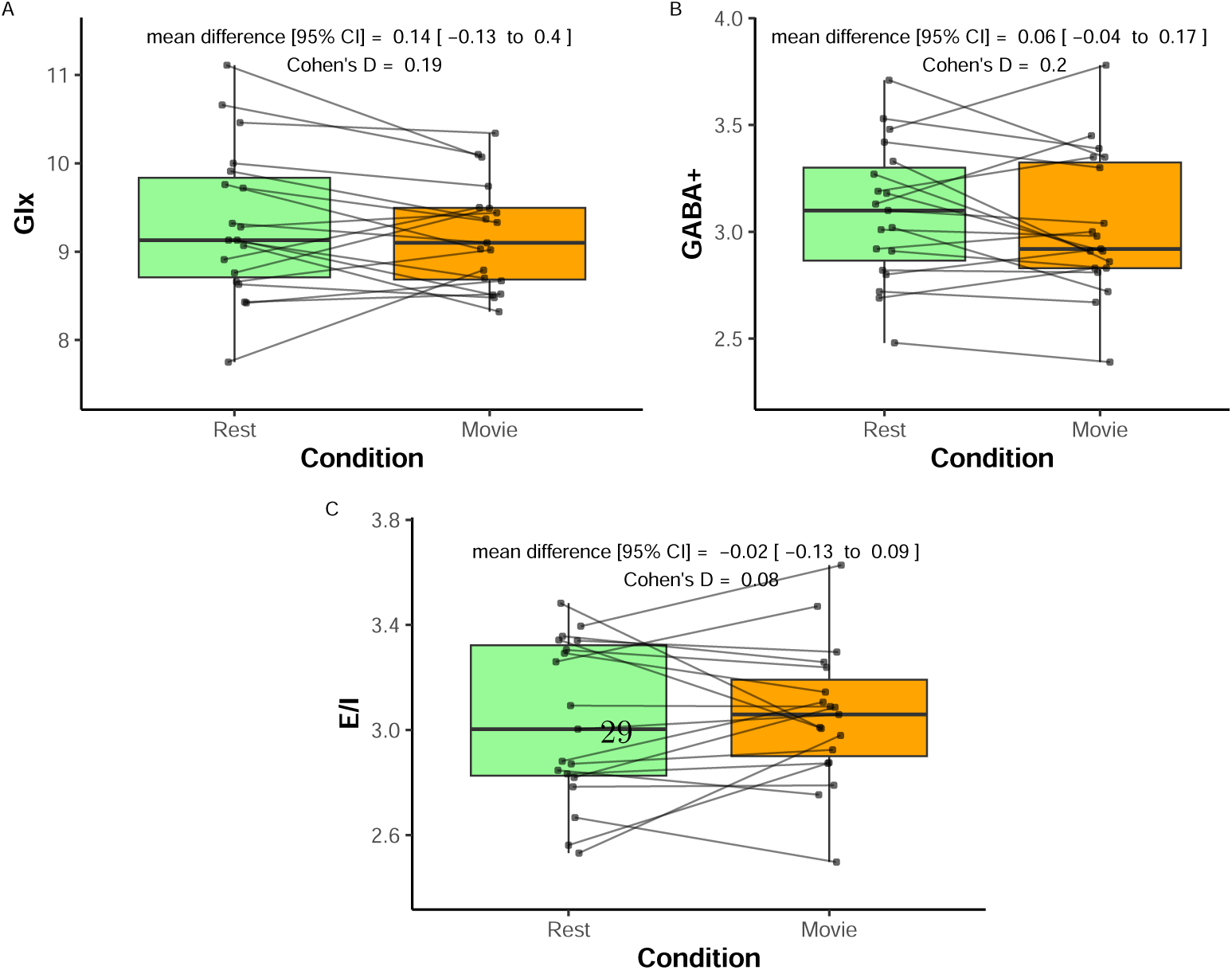
Paired comparison of metabolite values analyzed using Gannet during rest (green) and movie (orange) conditions. A) Glx; B) GABA+; and C) E:I. Paired dots represent the same participant across conditions. Mean difference with 95% confidence intervals, and as Cohen’s D are reported at the top of each plot.

H was not found to correlate with Glx, GABA+, or E:I, during rest or movie (Figure 9).

**Figure 9.**
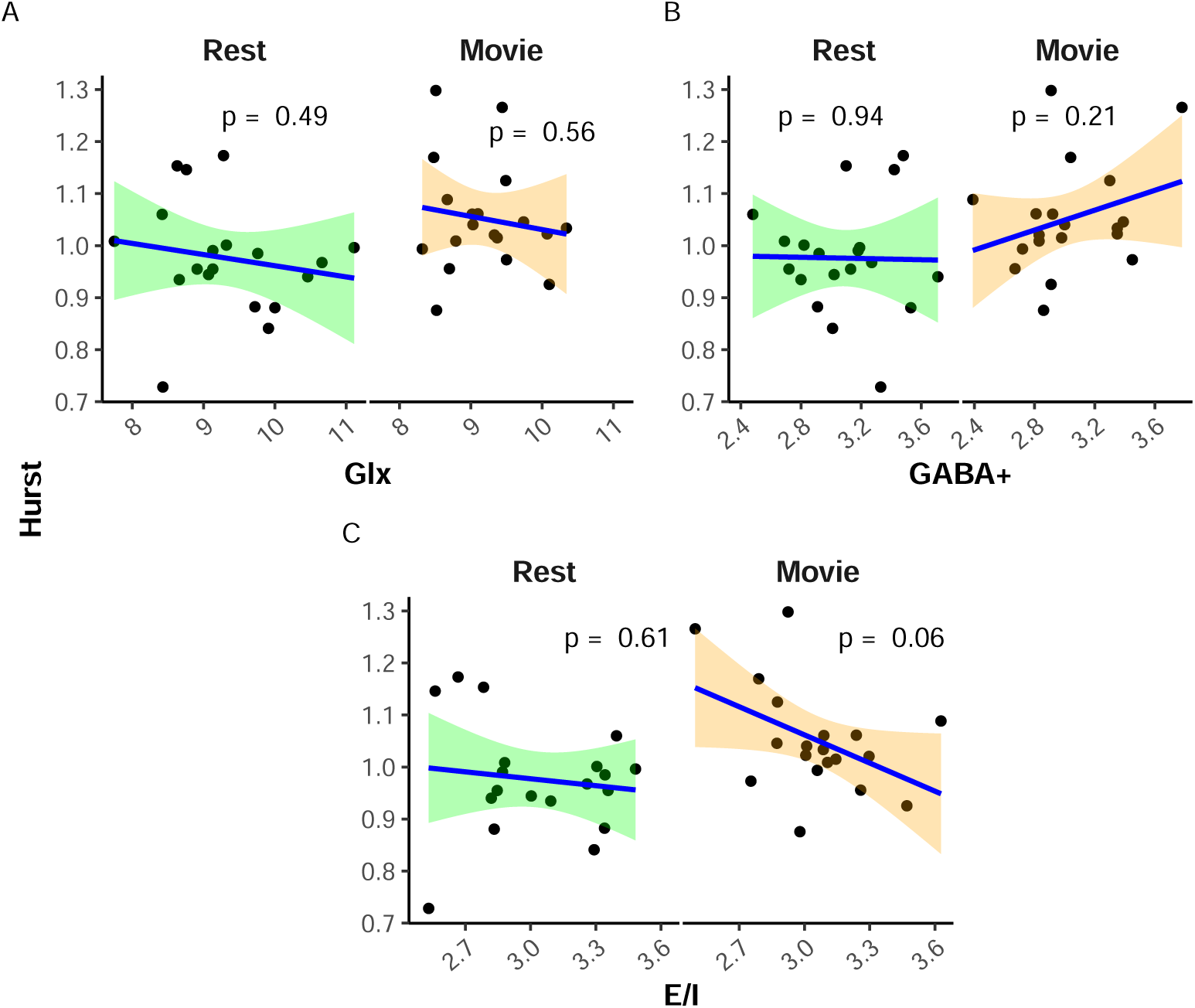
Scatter plots of H vs. metabolites when analyzed using Gannet. A) Glx; B) GABA+; and C) E:I. Rest is in green, while movie is in orange. p-values are reported at the top of each plot.

## Notes

https://github.com/WeberLab/EI_Hurst_Analysis

https://weberlab.github.io/EI_Hurst_Manuscript/

### Competing Interest Statement

The authors have declared no competing interest.

